# The histone modification regulator, SIN3, plays a role in the cellular response to changes in glycolytic flux

**DOI:** 10.1101/2025.01.15.633193

**Authors:** Imad Soukar, Anindita Mitra, Linh Vo, Melanie Rofoo, Miriam L. Greenberg, Lori A. Pile

**Affiliations:** Department of Biological Sciences, Wayne State University, Detroit, MI 48202, USA

**Keywords:** histone modification, respiration, gene regulation, energy metabolism, Drosophila

## Abstract

Epigenetic regulation and metabolism are connected. Epigenetic regulators, like the SIN3 complex, affect the expression of a wide range of genes, including those encoding metabolic enzymes essential for central carbon metabolism. The idea that epigenetic modifiers can sense and respond to metabolic flux by regulating gene expression has long been proposed. In support of this cross-talk, we provide data linking SIN3 regulatory action on a subset of metabolic genes with the cellular response to changes in metabolic flux. Furthermore, we show that loss of SIN3 is linked to decreases in mitochondrial respiration and the cellular response to mitochondrial and glycolytic stress. Data presented here provide evidence that SIN3 is important for the cellular response to metabolic flux change.

## 1. Introduction

Epigenetic regulation involves the post translational modification of N-terminal histone tails, which act as signaling molecules to recruit gene activators and repressors [1]. The most well-studied histone modifications include histone acetylation and methylation. Histone acetylation uses acetyl-CoA as the donor metabolite while histone methylation uses *S*-adenosylmethionine (SAM) as the donor metabolite. Acetyl-CoA is a key metabolic intermediate in central carbon metabolism, with its levels affected by the amount of available glucose [2]. SAM is produced by the one-carbon metabolism pathway, and levels of methionine in cell culture media correlate with intracellular SAM levels [3]. Perturbing pathways that generate donor metabolites influences histone modification levels. Disruption of acetyl-CoA generation leads to changes in histone acetylation levels [2,4–6]. Likewise, perturbation of SAM generating pathways affects histone methylation levels in Drosophila, yeast and mammalian cells [7–10].

Metabolic processes affect histone modifications and thus gene expression. Whether epigenetic regulators are essential for the cellular response to changes in metabolic flux, however, is not fully understood. Carbon flux through glycolysis is dependent on the availability of glucose. In budding yeast, glucose starvation leads to a reduction of acetyl-CoA concentration and global histone H3 acetylation levels [11]. Histone H3 lysine 9 acetylation (H3K9ac) is lost at the transcription start site (TSS) of many genes required for growth. While global H3K9ac levels decrease, a subset of genes exhibits an increase in H3K9ac upon glucose starvation. Deletion of either the Gcn5p lysine histone acetylase (KAT) or the Rpd3p histone deacetylase (HDAC) affects gene expression and histone modification response to altered glycolytic flux. The purpose of the studies presented here is to probe the relationship between metabolic flux and epigenetic regulators.

One of the earliest identified epigenetic regulating complexes is the SIN3 complex. *Sin3A* is conserved from yeast to humans [12]. *Sin3A* is an essential gene in *Drosophila melanogaster* and mice, wherein null mutation in *Sin3A* leads to embryonic lethality in mouse and Drosophila [13,14]. Additionally, the depletion of SIN3 levels leads to a significant decrease in cellular proliferation in Drosophila S2 cultured cells [15]. Furthermore, studies from the last two decades provide evidence that SIN3 is central to formation of multi-subunit complexes having many conserved functions [16]. As a scaffold protein, SIN3 interacts with histone modifying enzymes including HDAC1 (an HDAC) and KDM5 (a histone demethylase) [17,18]. The SIN3 complex regulates the post translational modification profiles of histone tails, thereby impacting the regulation of gene expression [19]. The SIN3 complex is often regarded as a repressor complex due to the presence of HDAC1, whose activity leads to histone deacetylation, often resulting in gene repression [20]. Genome-wide transcriptome studies, however, reveal that loss or reduction of SIN3 results in both gene repression and activation, suggesting that SIN3 complex activity is more complicated than simply through control of histone deacetylation [16]. While SIN3 is linked to regulation of genes that fall into a wide-ranging list of GO categories, genes linked to metabolic pathways are consistently identified in genome-wide studies, indicating that SIN3 from organisms including yeast, flies and mice is a key factor necessary for cellular homeostasis [21–25].

Consistent with the gene expression studies, our group and others have demonstrated that SIN3 is required for multiple cellular metabolic processes. For example, in Drosophila cultured cells, some of the genes repressed by the SIN3 complex are involved in central carbon metabolism, whereby the depletion of SIN3 results in elevated expression levels of numerous genes involved in glycolysis and the tricarboxylic acid cycle (TCA cycle) [24]. Analysis of mitochondrial function revealed that SIN3 impacts mitochondrial respiration, indicating that SIN3 regulation of metabolic genes affects cellular respiration under certain starvation conditions [26]. To study the relationship between SIN3 and bioenergetic regulation, our group performed a metabolomic analysis following *Sin3A* knockdown in Drosophila S2 cells [27]. This analysis revealed that SIN3 regulates the concentration of many metabolites associated with glycolysis and the TCA cycle. Additionally, deletion of SIN3 in yeast leads to low ATP levels and lack of growth in media requiring cellular respiration [26]. Together, the gene expression and metabolic studies indicate that SIN3 plays a crucial role in regulating cellular bioenergetics.

While reduction of SIN3 levels affects control of bioenergetic processes, it is unknown whether SIN3 is required for a cellular response to changes in metabolic homeostasis. Here, we test the hypothesis that SIN3 is essential for the ability of cells to respond to changes in glycolytic flux. We asked if alteration of metabolic flux affects the SIN3 complex, and its ability to regulate central carbon metabolism genes and cellular bioenergetics. We found that SIN3 regulation of metabolic genes is affected by alterations to glycolytic flux. We also demonstrate that SIN3 is required for the cellular response to mitochondrial and glycolytic stress. Furthermore, we observed that SIN3 interacts with a wide range of accessory proteins, which may contribute to the diversity of processes regulated by SIN3. These findings demonstrate the cross-talk between metabolism and the SIN3 epigenetic regulator. Additionally, these data indicate that SIN3 is required for the cellular response to changes in glycolytic activity.

## 2. Materials and Methods

### 2.1 Cell culture

Drosophila Schneider cell line 2 (S2) was cultured at 27°C in Schneider’s Drosophila Medium (ThermoFisher) containing 10% heat inactivated fetal bovine serum (Gibco) and 50 mg/ml gentamicin (Gibco). S2 cells carrying an HA-SIN3 transgene, used for the co-immunoprecipitation experiment, were cultured at 27°C in Schneider’s Drosophila Medium (ThermoFisher) containing 10% heat inactivated fetal bovine serum (Gibco), 0.1 mg/ml gentamicin (Gibco), and 0.1 mg/ml penicillin/streptomycin (Gibco). Generation of this cell line is described in Spain et al. 2010. Low glucose media was made by adding 0.6 g/L of calcium chloride, 0.4 g/L of sodium bicarbonate, 0.6 g/L of arginine, 2 g/L of trehalose, and either 11 mM or 2.75 mM glucose to Schneider’s Drosophila Medium from Caisson Labs (SCP03).

### 2.2 dsRNA production and RNA interference

The protocol for generating dsRNA against *Sin3A* and *GFP*, and RNA interference protocol is as previously described [15,28].

### 2.3 Western blot analysis

12 μg of protein extracts was used to analyze SIN3 and Tubulin protein levels. The protein concentration was determined by the DC assay from Bio-Rad, using the manufacturer’s protocol. Proteins were separated on 8% SDS-PAGE and transferred to a PVDF membrane (ThermoFisher Scientific) overnight. The membrane was then blocked for one hour with 5% milk made in PBS buffer with 0.2% Tween 20. Blots were incubated in primary antibody solution made with 5% milk, for two hours with a dilution of 1:1000 for SIN3 [29] and 1:500 for Tubulin (E7, DSHB). Secondary antibody incubation was done for one hour with a dilution of 1:3000. Western blot signal was detected using ECL prime (Cytiva Life Sciences). Western blot was done for all biological replicates to confirm *Sin3A* knockdown.

### 2.4 Glucose oxidase (GO) assay

Six million cells were treated with 1 mM, 5 mM or 20 mM concentrations of 2-Deoxy-D-glucose (2-DG) (Sigma-Aldrich) for six and 24 hours. Media was harvested and analyzed using the protocol described by the manufacturer (ab138884, Abcam).

### 2.5 Proliferation assay

Drosophila S2 cells were counted after 16 hours of 2-DG treatment using trypan blue (Lonza Biosciences).

### 2.6 Real-time quantitative PCR (qRT-PCR)

Total RNA was extracted from cells and converted to complementary DNA using the ImProm-II Reverse Transcription System (Promega). qRT-PCR reactions were carried out using PowerUp SYBR Green Master Mix (Applied Biosystems). The analysis was done using QuantStudio 3 Real-Time PCR system (ThermoFisher Scientific). Actin was used as the internal control for the gene expression studies. All primers used are listed in Table 1.

**Table 1:**
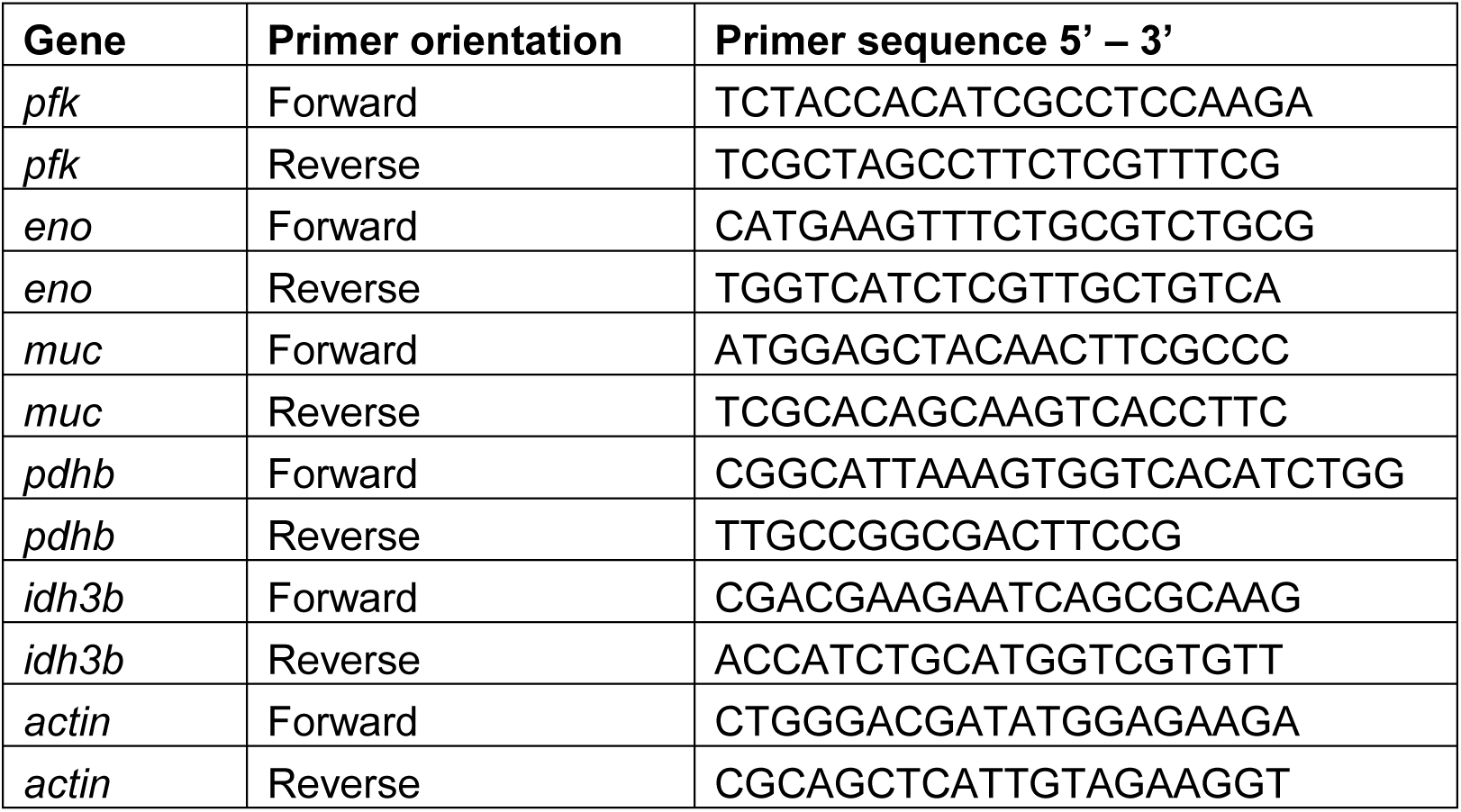
Primers used in this study.

### 2.7 Seahorse respiration assay

Drosophila S2 cells were seeded at 0.8-1.5 × 10^5^ cells/well in the XFe96 cell culture plate for 24 hours. Oxygen consumption rate was measured and analyzed using the Seahorse XF Cell Mito Stress Test Kit (103015-100) and the Seahorse XFe96 analyzer (Agilent). Basal respiration, maximal respiration, and coupling efficiency were calculated according to the manufacturer protocol. During the mitochondrial stress test, cells were treated with 2 μM oligomycin, 0.75 μM FCCP, and 0.5 μM Rotenone and Antimycin A. The experiment was done with eight technical and six biological replicates.

### 2.8 ATP assay

ATP levels from 10,000 cells were measured using the CellTitre-Glo 2.0 Cell Viability assay (Promega) following the manufacturer’s protocol. The SpectraMax i3x plate reader (Molecular Devices) was used to measure the amount of luminescence. The experiment was done with six technical and three biological replicates.

### 2.9 Seahorse glycolysis stress test

Drosophila S2 cells were seeded at 4 × 10^4^ cells/well for 24 hours, and the extracellular acidification rate was measured using the Seahorse XF Glycolysis Stress Test Kit (103020-100). The extracellular acidification rate was analyzed using the Seahorse XFe96 analyzer (Agilent). Basal glycolysis, glycolytic capacity, and glycolytic reserve were calculated as described by the manufacturer. Cells were treated with 11 mM glucose, 1 μM oligomycin and 50 mM 2-DG. The experiment was done with eight technical and three biological replicates.

### 2.10 Co-immunoprecipitation and analysis by TMT mass spec

#### 2.10.1 Sample Preparation

Protein samples in 25 μl SDS gel dissociation media were acidified by addition of 2 μl of 12% phosphoric acid then proteins were precipitated by addition of 200 μl 90% MeOH, 100 mM Triethylammonium bicarbonate buffer (TEAB, Honeywell Fluka cat# 60-044-974). Precipitation was initiated by incubating samples for one hour at 37°C and completed by overnight incubation at −20 C. Precipitates were pelleted by centrifugation for one minute at 16,000 x g, then washed by resuspending particulates in 400 μl of 90% MeOH, 10 mM TEAB. Precipitates were collected by centrifugation and dried on the bench then resuspended in 25 μl of 50 mM HEPES, pH 8.2. 0.25 μg Trypsin (Promega, V5113) was added to each sample and then incubated at 47°C for one hour then overnight at 37°C to complete the digestion. Following digestion, the samples were reduced and alkylated by incubating with 1 mM Dithiothretol (DTT, Sigma cat# D5545) for one hour at 37°C followed by addition of 3 mM Iodoacetamide (IAA, Sigma cat# I1149) for 30 minutes at room temperature in the dark followed by IAA quenching with addition of 1 mM DTT.

#### 2.10.2 Tandem Mass Tag (TMT) labeling and High pH-Reversed-Phase Peptide Fractionation

Samples were labeled with Tandem Mass Tag pro 16plex (TMTpro-16plex) reagents by addition of the selected reagent to each sample followed by incubation at room temperature for two hours. 140 µg of each TMTpro 16plex reagent (ThermoFisher, USA) was dissolved in 20 µL acetonitrile then immediately added to the peptide sample. The labeling reaction was allowed to proceed for two hours at room temperature. Each sample was evaluated for completeness of labeling then all samples were pooled and dried in a speed-vacuum, then fractionated using high pH reversed phase spin cartridges (ThermoFisher, USA). A step gradient of increasing ACN concentrations was applied to elute bound peptides in nine fractions. Each fraction was dried in a vacuum centrifuge and stored until analysis by mass spectrometry.

#### 2.10.3 Mass spectrometry analysis

Final analysis was performed on the samples using a Thermo Scientific Vanquish-Neo chromatography system with an Acclaim PepMap 100 trap column (100 µm × 2 cm, C18, 5 µm, 100Å), and Thermo Scientific Easy-Spray PepMap RSLC C18 75 μm x 25 cm column. A gradient starting at 6.8% acetonitrile and finishing at 30% acetonitrile 102 minutes later was used for all fractions. LC-MS/MS was performed on an Orbitrap Eclipse MS system operated with FAIMS (CV = −40, −55 and −70) and Real Time Search active. MS1 spectra were acquired at 120,000 resolution in the 400 to 1600 Da mass range with an AGC of 1e6. MS2 spectra were acquired in the ion trap and MS3 spectra triggered by the RTS were collected in the Orbitrap at 50 K resolution with a 100 to 500 Da window using Synchronized Precursor Selection (SPS) with 20 notches. Fragmentation for MS3 spectra was by HCD with a Collision Energy of 55, maximum injection time of 200 msec and an AGC target of 1^e^5.

### 2.11 Protein identification and quantification

Mass spectrometry data were processed by Proteome Discoverer version 2.4 using the Sequest HT algorithm with Percolator and the *Drosophila melanogaster* Uniprot FASTA database (downloaded March 25, 2021, 3,582 entries). The search parameters included trypsin with single missed cleavage. Variable modification by oxidation of M, and of protein N-termini by Acetylation, Met Loss or both Acetylation and Met loss. Carbamidomethylation of cysteine and TMTpro modification of lysine and of peptide N-termini were set as static modifications. Precursor and fragment mass tolerance were set to 10 ppm and 0.6 Da, respectively for MS2 spectra. For the entire data set, false discovery rate (FDR) was calculated by enabling the peptide sequence analysis using a decoy database, and a cut-off of 1% was used for identifications.

## 3. Results

### 3.1 SIN3 and inhibition of glycolysis regulate cell proliferation

Reduction of SIN3 results in altered expression of genes encoding glycolytic and TCA cycle enzymes as well as the concentration of many metabolic intermediates in these pathways [27]. Here, we wished to test the hypothesis that SIN3 gene regulatory activity is required for the cell to respond to changes in flux through glycolysis. To test this hypothesis, we perturbed glycolytic flux in control and *Sin3A* knockdown *Drosophila melanogaster* S2 cells by addition of 2-Deoxy-D-glucose (2-DG). 2-DG is a glucose analog that inhibits glycolysis by competitively binding hexokinase (Aft et al., 2002). The cellular uptake of 2-DG occurs via facilitated diffusion, followed by its conversion to 2-Deoxy-D-glucose-PO_4_ (2-DG-PO_4_) by hexokinase. Importantly, 2-DG-PO_4_ accumulation allosterically inhibits hexokinase [30]. To verify the inhibitory effects of 2-DG on glycolytic flux in Drosophila S2 cells, we measured the extracellular glucose levels of cells treated with increasing concentration of 2-DG or PBS vehicle using a glucose oxidation (GO) assay. As expected, treatment of cells with increasing concentrations of 2-DG led to the accumulation of extracellular glucose (Figure 1 A). Moreover, this effect of 2-DG on extracellular glucose persisted for 24 hours (Figure 1 A), indicating a sustained reduction in glycolytic rate due to 2-DG treatment.

**Figure 1.**
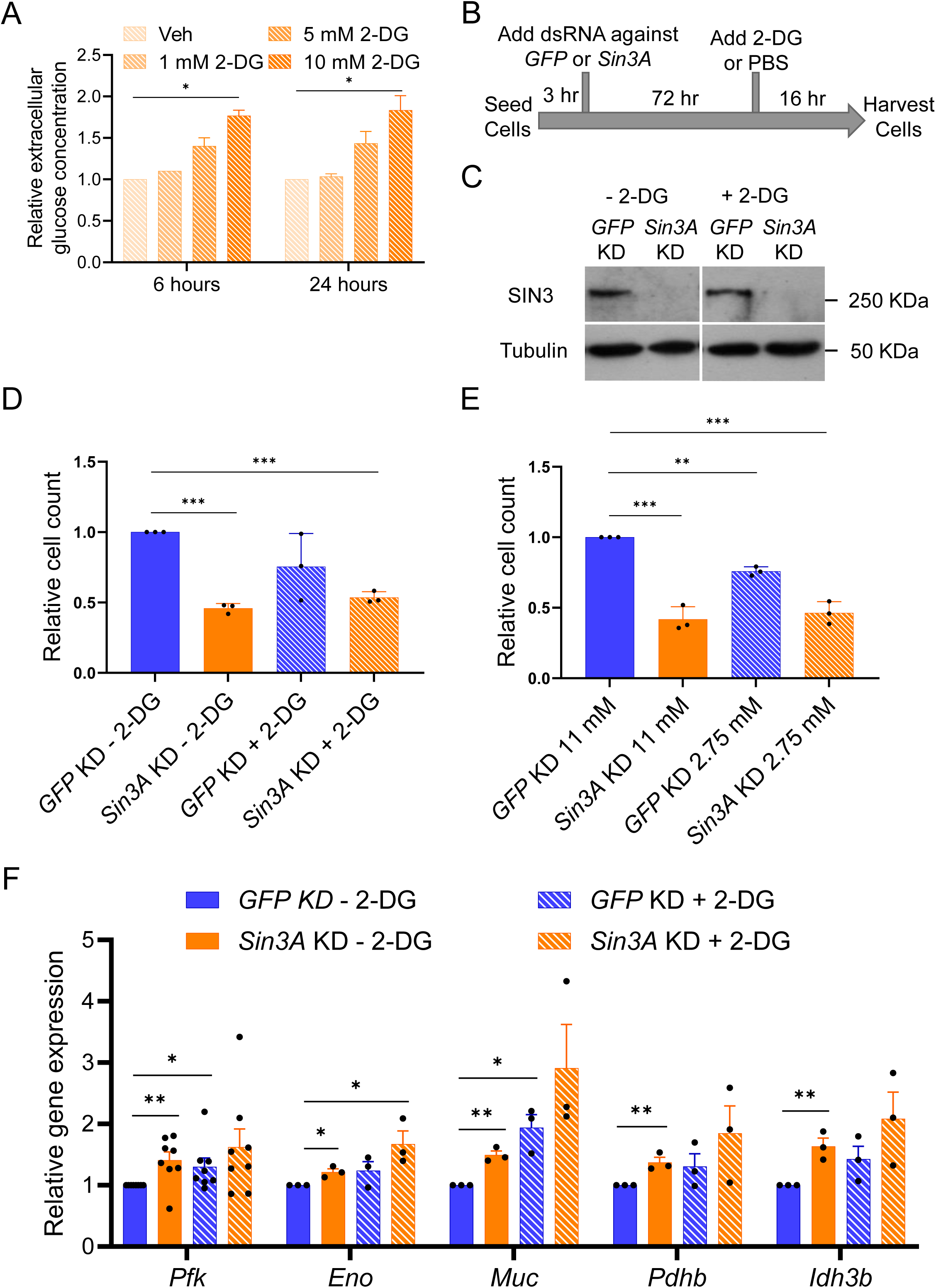
*Sin3A* knockdown leads to a proliferation defect and activation of metabolic genes. A. Extracellular glucose levels were measured after addition of PBS or treatment with increasing concentrations of 2-DG for six and 24 hours. B. Drosophila S2 cells were seeded for three hours and grown for an additional 72 hours after adding dsRNA against *GFP* or *Sin3A*. After 72 hours, cells were then treated with 1X PBS or 10 mM 2-DG for 16 hours before harvesting. C. SIN3 protein levels after knockdown in S2 cells were measured by SDS-PAGE followed by Western blotting. D-E. Cell proliferation of *Sin3A* knockdown cells was measured after growing cells in (D) 11 mM or 2.75 mM glucose or (E) 10 mM 2-DG for 16 hours. Three biological replicates were performed. F. qRT-PCR was used to measure the expression level of genes indicated. 10 mM 2-DG treatment was done for 16 hours. Three biological replicates were done for the genes except for *pfk*, where eight biological replicates were done. Error bars represent standard error of the mean. p-value * p<0.05, ** p<0.01, *** p< 0.001. The statistical test used was the two-tailed Student’s *t* test.

To examine the relationship between glycolytic flux and SIN3 regulatory function, we asked if the effect of SIN3 on cellular proliferation is affected by changes in glycolytic flux. To do this, we used RNA interference (RNAi) to knock down *Sin3A* or *GFP* followed by distinct treatments, including the addition of PBS or 2-DG (Figure 1 B), as well as culturing the cells in normal or low glucose media. Since Drosophila S2 cultured cells do not naturally express GFP, the knockdown of *GFP* serves as a control that activates the RNA interference machinery without affecting gene expression. To confirm the knockdown of *Sin3A*, we analyzed the protein levels by western blotting and observed a considerable reduction in SIN3 levels (Figure 1 C). Treatment with 2-DG did not affect the level of SIN3. Consistent with a previous report from our group [15], we observed a significant decrease in cell proliferation when SIN3 levels were reduced (Figure 1 D and E). Additionally, treatment of control cells with 2-DG for 16 hours resulted in a small decrease in proliferation relative to the control sample, although this change was not statistically significant (Figure 1 D). Interestingly, growing S2 cells in low glucose media (2.75 mM) for 16 hours led to a statistically significant decrease in cell proliferation (Figure 1 E). Combining *Sin3A* knockdown with 2-DG treatment or low glucose did not produce an additive effect on proliferation as cell counts were similar to those in the *Sin3A* knockdown only samples (Figure 1 D and E).

### 3.2 Some SIN3-regulated metabolic genes are sensitive to changes in metabolic flux

Changes to glycolytic flux have been previously shown to downregulate and upregulate the expression level of genes involved in glycolysis in mammalian regulatory T cells [31,32]. With 2-DG treatment, *hexokinase* expression level increases while *phosphofructokinase* expression level is reduced. To investigate whether changes to glycolytic flux similarly affect gene expression in Drosophila cells and to determine if SIN3 is required for the predicted gene expression response, we explored whether *Sin3A* knockdown combined with 2-DG treatment could result in the derepression of metabolic genes. To focus our study and identify potential interactions between SIN3 regulation and glycolytic flux, we concentrated on genes previously established as direct targets of SIN3 in comprehensive genome-wide studies. SIN3 binds the promoters of *phosphofructokinase* (*pfk)*, *enolase* (*eno*), *pyruvate dehydrogenase* (*pdhb*), *midline uncoordinated* (*muc*), and *isocitrate dehydrogenase* (*idh3b*) and the expression levels of each is upregulated following *Sin3A* knockdown in Drosophila S2 cells [24,33]. Notably, PFK plays a crucial role as one of the rate-limiting enzymes in glycolysis, while Eno facilitates the conversion of 2-phosphoglycerate to phosphoenolpyruvate. Pdhb and MUC are integral components of the Drosophila pyruvate dehydrogenase complex, responsible for generating acetyl-CoA that enters the TCA cycle. Idh3b generates alpha-ketoglutarate in the mitochondria, a metabolite essential for the activity of the JmjC family of histone demethylases, which are present within the SIN3 complex [34]. To measure gene expression, we performed RNA extraction and employed reverse transcription quantitative PCR (qPCR) analysis. Our results confirmed that SIN3 functions as a repressor of *pfk*, *eno*, *muc*, *pdhb* and *idh3b* genes (Figure 1 F), in line with our previous findings (Pile et al., 2003). Addition of 2-DG to cells with wild-type levels of SIN3 resulted in a statistically significant increase in the expression of *pfk* and *muc* genes (Figure 1 F). To investigate the interplay between SIN3 regulation and changes in glycolytic flux, we examined gene expression changes in cells treated with 2-DG while simultaneously depleting SIN3 levels. All tested genes showed elevated expression, comparable to that observed in cells treated having either knockdown of *Sin3A* or following 2-DG treatment. With the exception of *eno*, these changes comparing the individual treatments to the dual treatment, however, were not statistically significant (Figure 1 F). Notably, an additive effect was not observed at any of the tested genes, indicating that the effect of 2-DG is redundant in the context of *Sin3A* knockdown. These findings suggest that SIN3 typically acts as a repressor for certain central carbon metabolism genes and a decrease in glycolytic flux affects their expression. Overall, these data predict that SIN3 regulatory function is involved in the cellular response to changes in glycolytic flux.

### 3.3 SIN3 is required for the ability of cells to respond to metabolic stress

ATP serves as the primary energy source within cells and is a key byproduct of mitochondrial respiration [35]. SIN3 regulates genes encoding glycolytic and TCA cycle enzymes and impacts levels of metabolites necessary for ATP production [27]. Here, we aimed to investigate whether SIN3 is required for the cellular bioenergetic response to changes in metabolic flux. For our study, we knocked down *GFP* or *Sin3A* in S2 cells, followed by treatment with either PBS or 2-DG. Following this treatment, we measured the oxygen consumption rates of cells using the Seahorse XFe96 analyzer (Agilent) (Figure 2 A). In addition to assessing the basal respiration rates (Figure 2 B), we also evaluated the maximal respiration and coupling efficiency (Figure 2 C and D). Basal respiration represents the energy demand of cells under normal conditions. We observed a significant reduction in basal respiration upon the knockdown of *Sin3A*. Similarly, the addition of 2-DG to cells resulted in a comparable decrease in basal respiration (Figure 2 B). This phenotype was also observed in cells with reduced SIN3 levels that were also treated with 2-DG (Figure 2 B). Maximal respiration, which provides insight into the ability of the cell to respond to metabolic stress [36], exhibited a decrease when SIN3 levels were perturbed. Treatment with 2-DG alone did not induce a significant change in maximal respiration when compared to PBS-treated *GFP* knockdown cells. Maximal respiration in 2-DG treated cells was notably higher compared to both untreated and treated *Sin3A* knockdown cells (Figure 2 C). The reduction in maximal respiration observed in *Sin3A* knockdown cells suggests that SIN3 plays a role in the cellular capacity to respond to energy demands. Lastly, we measured coupling efficiency, which quantifies the extent to which mitochondrial respiration is linked to ATP production [37]. The reduction in SIN3 levels led to a decrease in coupling efficiency for both untreated and 2-DG treated cells. Furthermore, the addition of 2-DG alone resulted in a small but significant reduction in coupling efficiency (Figure 2 D). These findings reinforce the idea that SIN3 is essential for maintaining mitochondrial function and enables cellular responses to energy demands.

**Figure 2.**
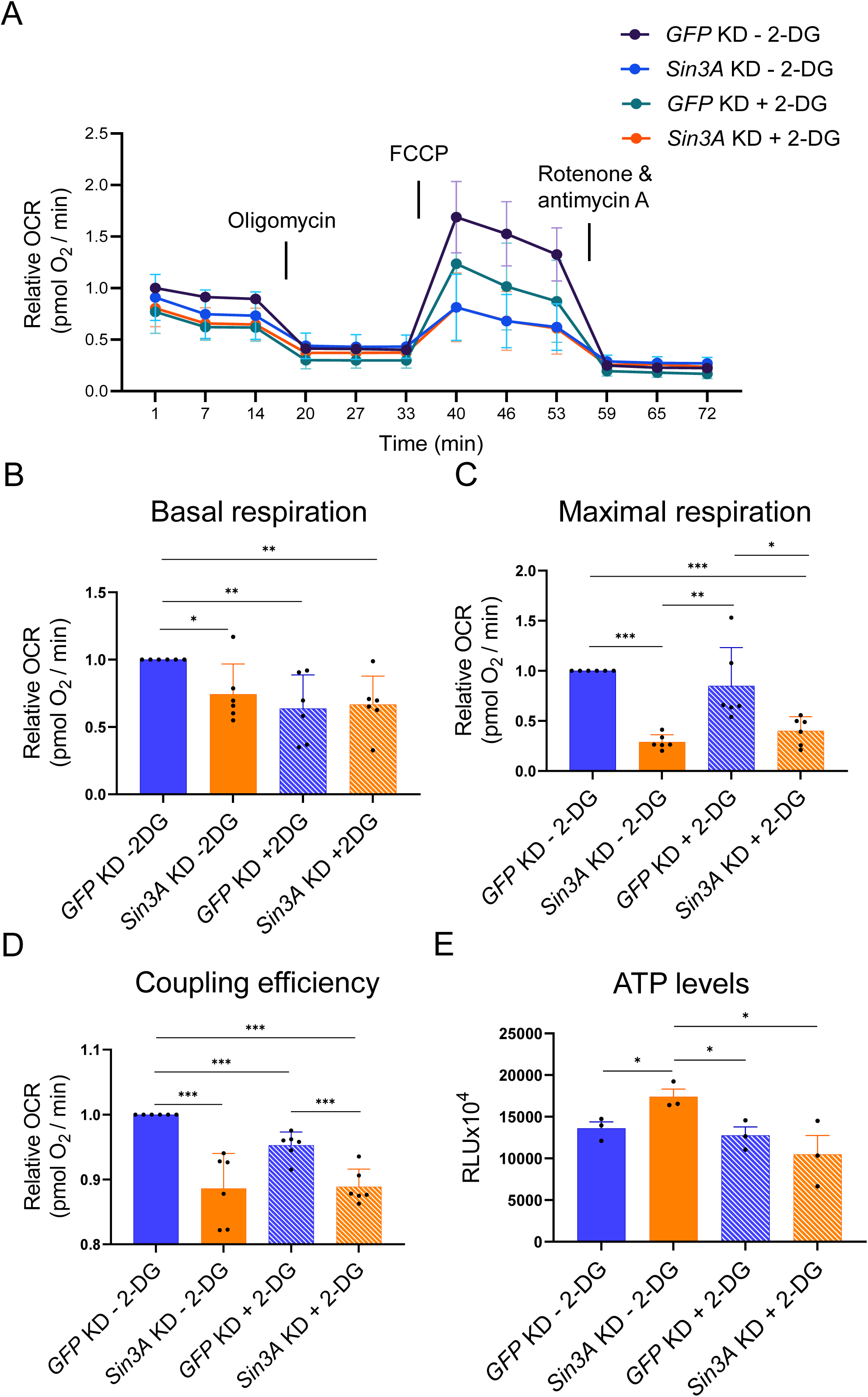
SIN3 regulates mitochondrial respiration and ATP levels. A. Mitochondrial stress test profile of *GFP* or *Sin3A* knockdown cells treated with PBS or 2-DG. B-D. Basal respiration rates (B), maximal respiration (C), and coupling efficiency (D) were calculated from (A). E. ATP levels were calculated from *GFP* or *Sin3A* knockdown cells treated with PBS or 2-DG. Six biological replicates were conducted for (A-D) and three biological replicates were done for (E). Error bars represent standard error of the mean. p-value * p<0.05, ** p<0.01, *** p< 0.001. The statistical test used was the two-tailed Student’s *t* test.

To gain further insights into the role of SIN3 in respiration and energy response, we examined whether the impact of SIN3 on ATP levels is affected by 2-DG treatment. Previous studies conducted by our group demonstrated that *Sin3A* knockdown could influence ATP levels under certain starvation conditions [26]. To measure ATP levels, we utilized a luminescence-based assay in cells with knockdown of *GFP* or *Sin3A*, which were treated with either PBS or 2-DG. Interestingly, *Sin3A* knockdown led to a significant increase in ATP levels compared to *GFP* knockdown control cells (Figure 2 E). This increase in ATP levels, however, was not observed in *Sin3A* knockdown cells treated with 2-DG. Moreover, the addition of 2-DG to *GFP* knockdown cells did not result in a significant change compared to non-treated control cells (Figure 2 E). These results indicate that SIN3 activity impacts ATP levels under normal metabolic conditions. When glycolysis is inhibited, the need for SIN3 is mitigated or compensated through other mechanisms as loss has no effect on ATP levels under these conditions.

### 3.4 SIN3 does not regulate basal glycolysis but is required for a glycolytic stress response

The observation that knockdown of *Sin3A* leads to an increase in ATP levels in PBS-treated cells but not in 2-DG-treated cells prompted us to explore the ways in which SIN3 influences ATP levels under these distinct metabolic conditions. Interestingly, despite the reduction in mitochondrial respiration (Figure 2 B), *Sin3A* knockdown cells exhibit increased ATP levels. One possible explanation for this change in ATP concentration is that *Sin3A* knockdown enhances the glycolytic rate, which is then suppressed when glycolysis is inhibited by 2-DG. Another potential mechanism is that *Sin3A* knockdown cells exhibit reduced ATP consumption, leading to higher overall ATP levels compared to control cells. Consequently, when treated with 2-DG, which acts as an additional stressor, ATP levels decrease compared to *Sin3A* knockdown cells treated with PBS. To investigate these hypotheses, we measured the glycolytic rate of cells using the Seahorse XFe96 analyzer (Agilent) in both *GFP* and *Sin3A* knockdown cells cultured in media with either 11 mM or 2.75 mM glucose (Figure 3 A). Notably, we employed different glucose concentrations instead of 2-DG to modulate glycolytic flux, as 2-DG was used in the assay. Additionally, as part of the protocol, all cells, regardless of the treatment, were subjected to a one-hour starvation period before measuring glycolytic rates. Unexpectedly, we found that the basal glycolytic rate remained unaffected by changes in SIN3 levels or glucose concentration (Figure 3 B). Next, we aimed to determine if SIN3 regulates the cellular response to glycolytic stress. For this purpose, we measured the glycolytic capacity, which reflects the maximum rate at which a cell can undergo glycolysis [38]. Additionally, we assessed the glycolytic reserve, which quantifies the ability of the cells to respond to glycolytic stress [39]. Cells with high glycolytic capacity and reserves are better equipped to handle glycolytic stress compared to cells with low glycolytic capacity and reserve. The knockdown of *Sin3A* in cells cultured in both 11 mM and 2.75 mM glucose media led to a significant decrease in glycolytic capacity. In contrast, growing *GFP* knockdown control cells in 2.75 mM glucose media did not result in a significant change in glycolytic capacity (Figure 3 C). In contrast, growing *GFP* knockdown control cells in 2.75 mM glucose media did not result in a significant change in glycolytic capacity. Furthermore, *Sin3A* knockdown resulted in a reduction in glycolytic reserve, which was observed in *Sin3A* knockdown cells grown in 2.75 mM glucose media, compared to control cells grown in 11 mM glucose media (Figure 3 D). Cells grown in low glucose media but without perturbing SIN3 levels exhibited a non-significant increase in glycolytic capacity and reserve (Figure 3 C and D). These findings indicate that SIN3 does not regulate the basal glycolytic rate but does modulate the parameters involved in the cellular ability to respond to glycolytic stress. The basal glycolysis data indicate that the increase in ATP levels in *Sin3A* knockdown cells (Figure 2 E) might be due to the reduction in ATP consumption and not due to the upregulation of glycolysis.

**Figure 3.**
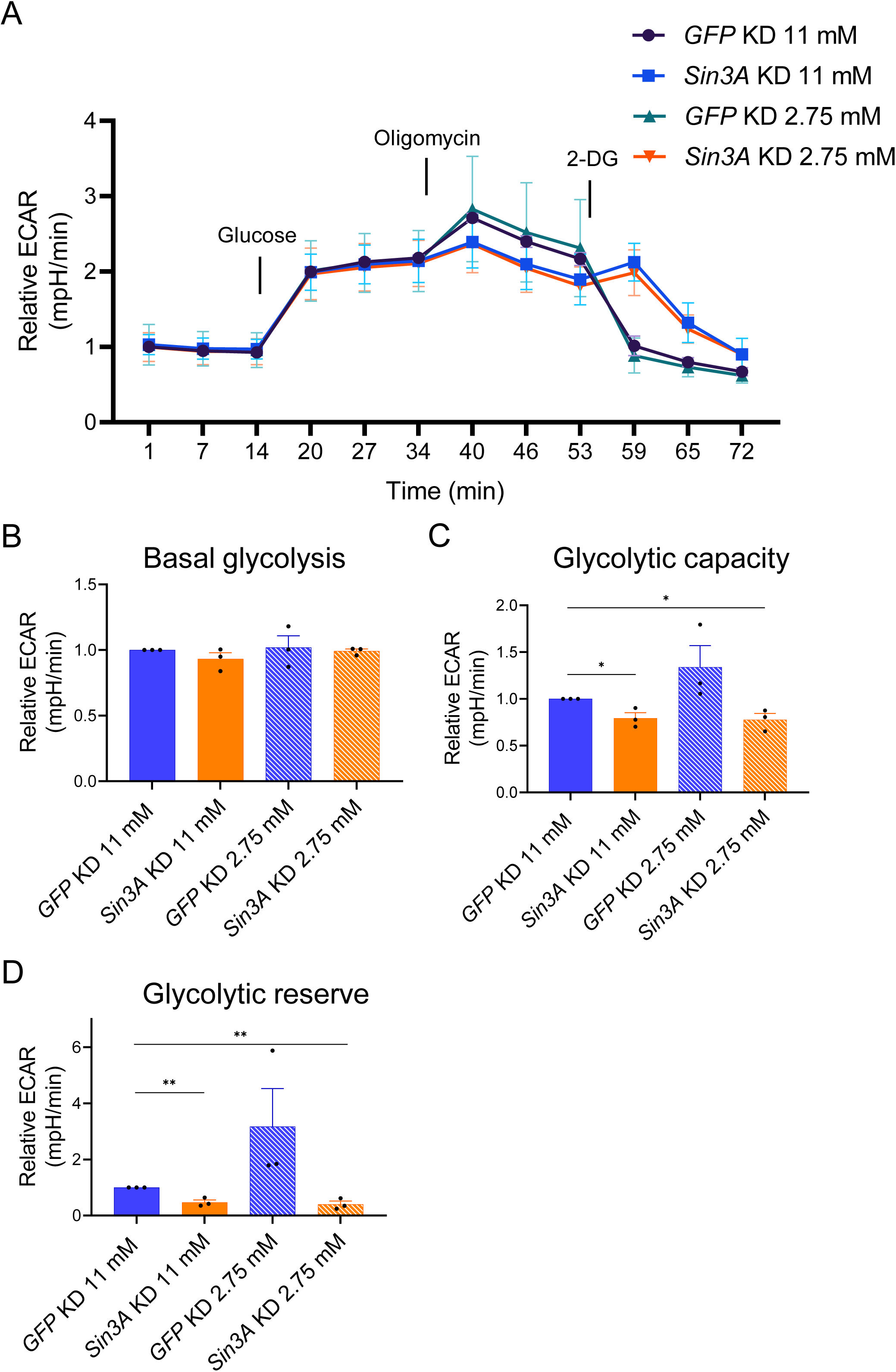
SIN3 regulates the response to glycolytic stress. A. Glycolysis stress test profile *GFP* or *Sin3A* knockdown cells grown in 11 mM or 2.75 mM glucose. B-D. Basal glycolysis rate (B), glycolytic capacity (C) and glycolytic reserve (D) were calculated from (A). Three biological replicates were performed. Error bars represent standard error of the mean. p-value * p<0.05, ** p<0.01, *** p< 0.001. The statistical test used was the two-tailed Student’s *t* test.

### 3.5 SIN3 complex composition is not affected by changes in glycolytic flux

Next, we aimed to investigate the way in which SIN3 regulates metabolic genes and influences the cellular response to changes in glycolytic flux. We asked whether the composition of the SIN3 complex changes under 2-DG treatment. We hypothesized that alterations in glycolytic flux are sensed by SIN3 and/or other complex components, leading to changes in the complex composition. To test this hypothesis, we used a cell line expressing ectopic SIN3 that has an HA tag [40]. We immunoprecipitated HA-tagged SIN3 and analyzed its interactors using tandem mass tag (TMT) multiplexing with LC-MS/MS that included Real Time Search (RTS) and Synchronous Precursor Selection (SPS). TMT multiplexing with RTS and SPS allows for efficient and quantitative analysis by labeled peptides [41], enabling us to simultaneously measure the abundance of SIN3 complex components isolated from cells treated with either PBS or 2-DG. The quantitation by mass spectrometry identified a large set of SIN3 complex interacting proteins. Our analysis of the complex under control and 2-DG conditions, however, revealed no changes in its composition (Figure 4 A). Under both conditions, we observe proteins that had previously been shown to be in the complex with SIN3 at similar abundance to SIN3 itself, while other proteins are present at sub-stoichiometric amounts.

**Figure 4.**
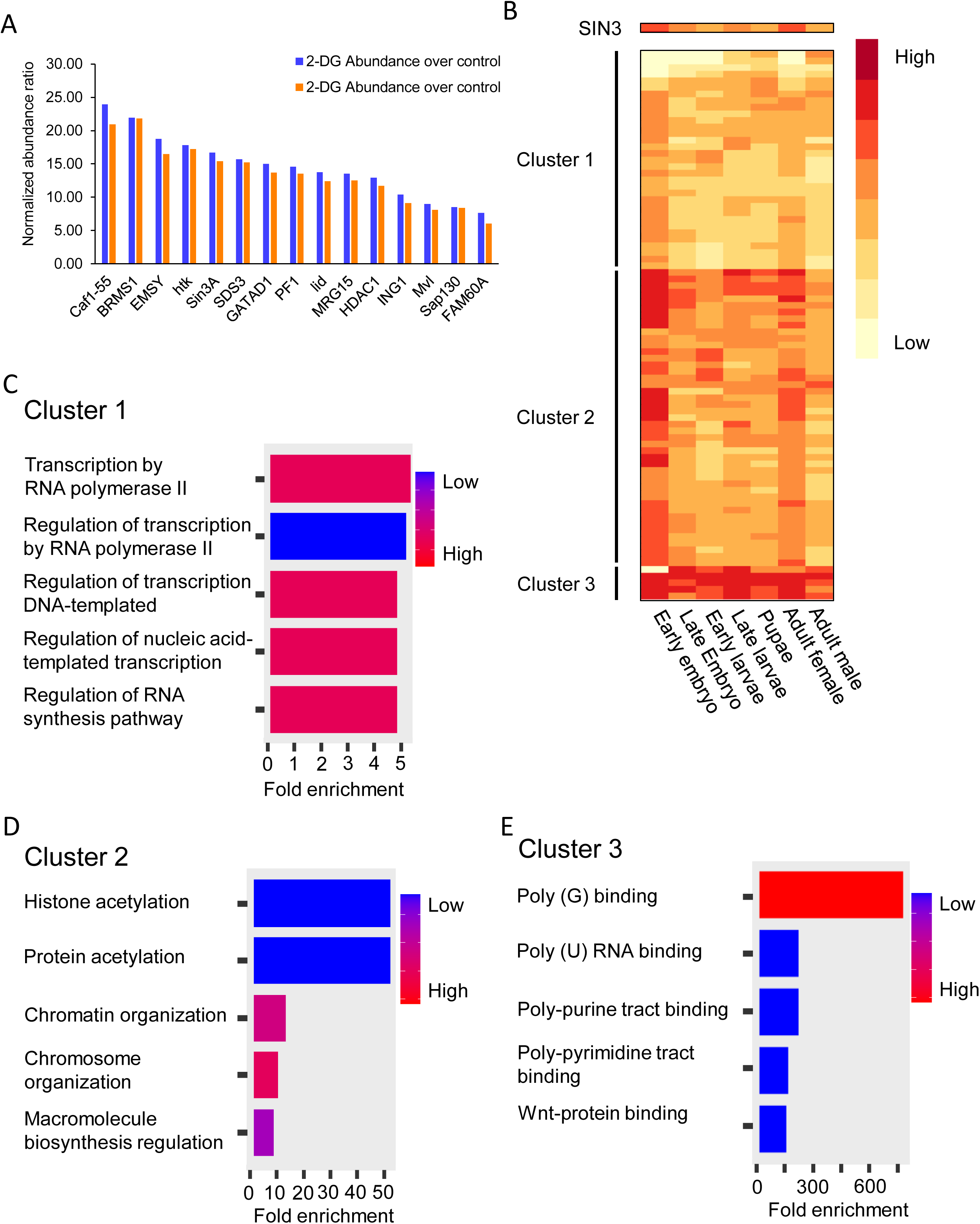
The SIN3 complex composition does not change when cells are treated with 2-DG. A. Relative abundance ratio is plotted for each of the proteins that were significantly enriched over the control and have similar abundance level to SIN3. Protein abundance was normalized to SIN3 abundance. B. 171 proteins that were statistically enriched in the SIN3 pulldown sample, compared to the no tag control, were clustered and visualized by the heat map. Five biological replicates were done. C-E. Gene ontology analysis was done by ShinyGO 0.77 using default settings. Clusters were from Figure 4B.

From the heterogeneous collection of SIN3 interactors, we next aimed to identify constitutive proteins that are most likely present in all SIN3 complexes as well as proteins that act as accessory complex components, interacting in a context dependent manner. To identify constitutive complex components, we employed a stringent analysis method. Firstly, to be considered a constitutive complex component, the proteins identified have to exhibit significant enrichment (P value < 0.05) compared to the no tag control set. Secondly, the protein abundance has to be similar to that of SIN3 enrichment. Our rationale is that constitutive complex components should be present in similar abundance to SIN3. Using these parameters, we successfully identified constitutive complex components, as previously reported by our group [40] (Figure 4 A). Additionally, we identified some proteins known to interact with mammalian SIN3, such as FAM60A, MRG15 and Sap30 [42,43], and found they also interact with the Drosophila SIN3 complex (Figure 4 A). FAM60A, MRG 15 and Sap30 were not discovered in our original SIN3 complexes, perhaps due to the different type of analysis compared to the current method using TMT multiplexing which minimizes missing values and allows deeper proteome coverage than standard data dependent analysis.

Using a less stringent analysis in which we included proteins significantly enriched compared to the no tag control samples, we identified a total of 171 proteins that exhibited significant enrichment in either treated or non-treated samples. This set included the 15 constitutive protein partners and 156 accessory factors. We predict that the accessory proteins bind transiently and might interact with SIN3 only in certain contexts. Among these proteins, two drew particular interest from our group: RNA Polymerase II subunit A (Rpb1) and Negative elongation factor A (NELF-A). Both proteins interact with SIN3 in treated and non-treated samples. In a recent bioinformatics report from our group, we found that SIN3 regulatory action correlates with Rpb1 and NELF-A binding [44]. The interaction between SIN3, Rpb1 and NELF-A supports our hypothesis that SIN3 regulates a subset of genes by binding to and regulating the elongation dynamics of Rpb1.

To complement the interaction studies, we analyzed the expression profile of the 171 proteins that exhibited significant enrichment in the SIN3-immunoprecipitated samples compared to the no tag control set. Given that SIN3 has been found to interact with some distinct proteins in different tissues [40], we investigated whether proteins identified in our analysis show a similar expression profile to SIN3 throughout Drosophila development. This type of analysis will support our hypothesis that some SIN3 interactors are constitutive while accessory interactors change in a context dependent manner. This analysis, however, is limited considering that the SIN3 complex in our analysis was immunoprecipitated from the S2 cell line, originally derived from late stage embryos [45]. We obtained gene expression data determined at distinct Drosophila developmental stages from Flybase.com [46] and analyzed the expression patterns of SIN3 interactors. Through k-means clustering, we identified three distinct clusters of gene expression patterns (Figure 4 B). We next performed gene ontology (GO) analysis on the three gene clusters using ShinyGO 0.77 [47,48]. Among the clusters, cluster 2 exhibited the highest similarity to the expression pattern of *Sin3A* (Figure 4 B). Cluster 2 contains proteins that are constitutive partners of the SIN3 complex and possibly work through the canonical mechanism through which SIN3 is known to regulate genes, specifically by modulating histone modifications (Figure 4 D). Cluster 1 includes proteins involved in RNA pol II elongation (Figure 4 C). These proteins might be a part of a more dynamic SIN3 complex throughout developmental stages. In other words, these proteins might be interacting with SIN3 in a transient fashion, and when their expression levels are high in some developmental stages, it is possible that they interact with SIN3 for a particular regulatory function. Lastly, cluster 3 consists of genes encoding proteins that have a high level of expression throughout development. GO analysis of cluster 3 indicates that these proteins play a role in RNA binding regulation (Figure 4 E). Collectively, these findings highlight potential modes of regulation through which different SIN3 complexes might influence the expression of a diverse set of genes.

## 4. Discussion

In this study, we analyzed regulatory links between the epigenetic regulator SIN3 and cellular bioenergetics. By perturbing the glycolytic flux through 2-DG treatment or by culturing cells with or without SIN3 in low glucose conditions, we investigated the cross-talk between SIN3 and metabolic flux. We note a reliance of cells on the SIN3 complex for responding to altered glycolytic activity. Our findings align with our previous work linking another SIN3 paralog to metabolic stress response [49]. We examined the SIN3 interactome and found that the complex is unaffected by 2-DG treatment. By analyzing the gene expression profiles of SIN3 interactors across development, we were able to distinguish core complex components from those that likely interact transiently.

Specifically, we observed that metabolic gene expression levels were higher in *Sin3A* knockdown cells compared to controls, resembling the pattern observed in wildtype cells treated with 2-DG (Figure 1 E). Furthermore, when *Sin3A* knockdown cells were subjected to 2-DG treatment, we observed a comparable increase in gene expression similar to that observed in control cells treated with 2-DG (Figure 1 C). This finding suggests that certain metabolic genes, typically repressed by SIN3, lose their repression in response to 2-DG treatment (Figure 7). Although the fold change in gene expression was modest, SIN3 is proposed to function as a soft repressor. This type of repression involves fine-tuning the expression of genes rather than exerting on/off control [50]. Additionally, we measured oxygen consumption rates and determined that SIN3 contributes to the cellular response to metabolic stress (Figure 3). Following stress of mitochondrial function through the addition of inhibitors and activators, we measured various bioenergetic parameters, including basal respiration (Figure 2 B) and maximal respiration (Figure 2 C), which measures the ability to respond to metabolic stress. Analysis of these data leads us to infer that the role of SIN3 in the regulation of mitochondrial function is essential for the ability of the cell to respond to metabolic stress due to inhibition of glycolysis.

Interestingly, ATP levels in *Sin3A* knockdown cells are higher compared to cells harboring wildtype levels of SIN3 (Figure 2 E). This was surprising since basal respiration rates were low in cells with reduced SIN3 (Figure 2 B). In contrast, upon reduction of SIN3 levels combined with growth in the presence of 2-DG, ATP levels decreased compared to wildtype (Figure 2 E), consistent with a previous finding from our group [26]In that study, Barnes et al., used depleted media by diluting Drosophila S2 media with PBS, while our study here used 2-DG to affect the glycolytic flux, treatments that ultimately yielded similar results. The increase in ATP levels observed when SIN3 levels are reduced can be attributed to two potential reasons. Firstly, it may be a result of increased glycolytic activity, leading to increased ATP production. Secondly, it could be due to a reduction in ATP consumption. Notably, cellular proliferation events, such as protein and DNA synthesis, rank among the highest consumers of ATP within the cell [51]. To determine the underlying reason, we conducted measurements of glycolytic rates in *Sin3A* knockdown cells (Figure 3 A). Interestingly, we found no significant difference in glycolytic rates compared to control cells, suggesting that the increase in ATP levels in the *Sin3A* knockdown cells is not due to an upregulation of glycolysis (Figure 3 B). Instead, it is more plausible that the elevated ATP levels result from a decrease in ATP consumption. This possibility is supported by the observation that *Sin3A* knockdown results in an inhibition of cellular proliferation (Figure 1 C and D), a major ATP-consuming process [51].

The composition of the SIN3 complex was essentially unchanged during altered glycolytic flux. The TMT mass spectrometry data did reveal that there are possibly multiple SIN3 complex assemblies that control gene expression. One such SIN3 complex assembly involves the regulation of RNA pol II elongation. This gene regulatory mechanism is supported by previous reports suggesting a connection between HDAC activity and RNA pol II pausing near the promoters of specific genes [52]. Consistent with this, our recent bioinformatics analysis demonstrates that SIN3 repressed genes exhibit increased binding of RNA pol II and the pausing factor NELF-A near the transcription start site, compared to genes activated by SIN3 [44]. Further analysis of the mass spectrometry data revealed a set of constitutive SIN3 complex components that share a similar expression pattern throughout Drosophila development. GO analysis of these genes is consistent with the canonical mechanism in which SIN3 regulates genes through histone deacetylation.

In summary, our findings indicate that SIN3 is required for the cellular response to changes in metabolic flux due to inhibition of glycolysis. Additionally, we determined that SIN3 plays a vital role in the cellular response to both mitochondrial and glycolytic stress. Based on our current understanding, we propose a working model wherein changes in glycolytic flux caused by 2-DG result in a reduction of oxygen consumption rate, which is detected by the SIN3 complex. Consequently, this sensing mechanism leads to the derepression of key metabolic genes (Figure 5).

**Figure 5.**
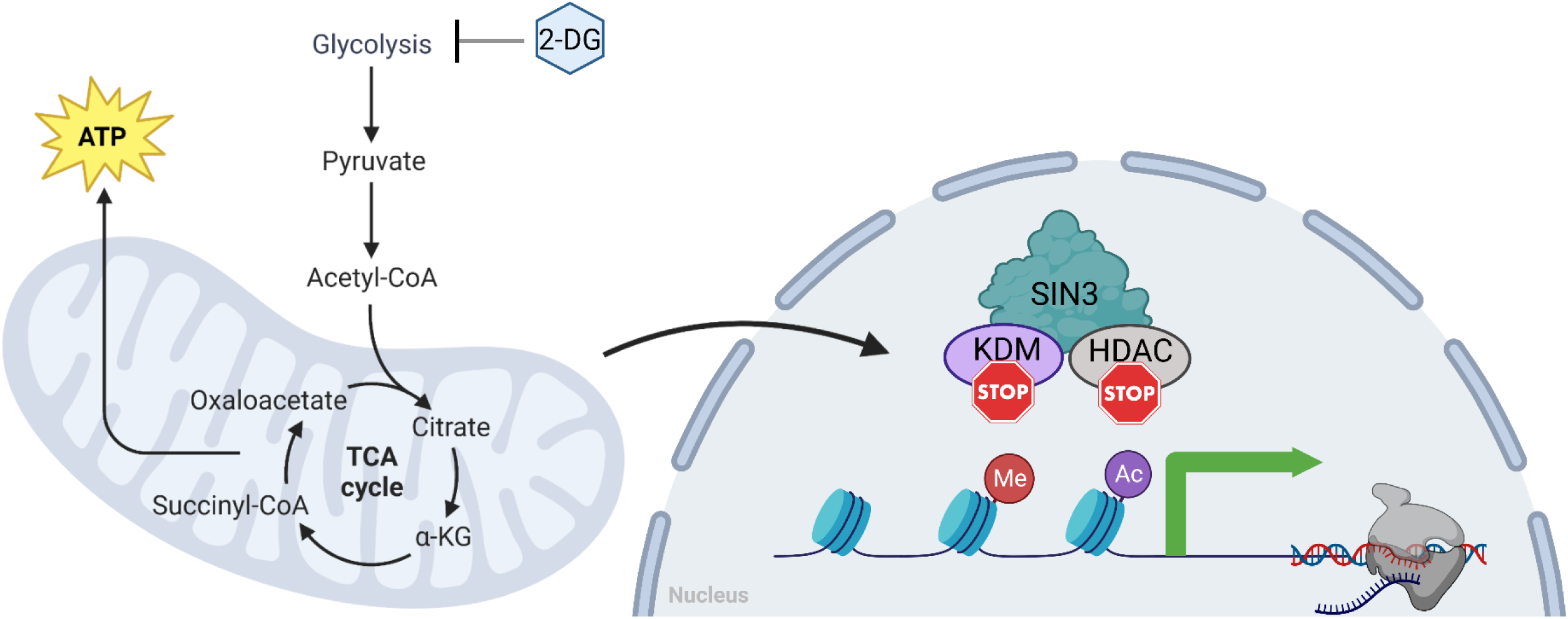
SIN3 is responsive to changes to the glycolytic flux. Our current model predicts that SIN3 acts as a repressor on numerous metabolic genes. In response to glycolytic flux change, mitochondrial respiration rates are affected, which are then sensed by SIN3. Consequently, SIN3 repression on those metabolic genes is lost, leading to an increase in their expression. This sensing mechanism by SIN3 is not due to changes in the composition of the complex, but rather likely due to changes in the activity of the complex. One possibility is that SIN3 is post translationally modified, allowing it to differentially regulate metabolic genes to affect the bioenergetics of cells.

## Glossary

SAM: *S*-adenosylmethionine
H3K9ac: Histone H3 lysine 9 acetylation
TSS: Transcription Start Site
KAT: lysine histone acetylase
HDAC: Histone Deacetylase
TCA cycle: Tricarboxylic Acid Cycle
2-DG: 2-Deoxy-D-glucose
RNAi: RNA interference
qPCR: quantitative PCR

## Data Availability

The mass spectrometry data can be accessed through the PRIDE repository with the accession number: PXD058616.

## Acknowledgments

We acknowledge the assistance of the Wayne State University Proteomics Core that is supported through NIH grants P30ES036084, P30CA022453 and S10OD030484.

## Conceptualization

L.A.P.; Investigation: I.S., A.M., L.V., M.R.;

Formal analysis: I.S., A.M., L.V., M.R.;

Writing - original draft: I.S., L.A.P.;

writing – review and editing: I.S., A.M., L.V., M.R., M.G., L.A.P.;

Supervision: M.G., L.A.P.

## References

[1] A.J. Bannister, T. Kouzarides, Regulation of chromatin by histone modifications, Cell Res, 21 (2011) 381–395.

[2] A.A. Cluntun, H. Huang, L. Dai, X. Liu, Y. Zhao, J.W. Locasale, The rate of glycolysis quantitatively mediates specific histone acetylation sites, Cancer Metab, 3 (2015) 10.

[3] J. Jelleschitz, Y. Zhang, T. Grune, W. Chen, Y. Zhao, M. Jia, Y. Wang, Z. Liu, A. Höhn, Methionine restriction - Association with redox homeostasis and implications on aging and diseases, Redox Biol, 57 (2022) 102464.

[4] A. Moussaieff, M. Rouleau, D. Kitsberg, M. Cohen, G. Levy, D. Barasch, A. Nemirovski, S. Shen-Orr, I. Laevsky, M. Amit, D. Bomze, B. Elena-Herrmann, T. Scherf, M. Nissim-Rafinia, S. Kempa, J. Itskovitz-Eldor, E. Meshorer, D. Aberdam, Y. Nahmias, Glycolysis-mediated changes in acetyl-CoA and histone acetylation control the early differentiation of embryonic stem cells, Cell Metab, 21 (2015) 392–402.

[5] H. Takahashi, J.M. McCaffery, R.A. Irizarry, J.D. Boeke, Nucleocytosolic acetyl-coenzyme a synthetase is required for histone acetylation and global transcription, Mol Cell, 23 (2006) 207– 217.

[6] K.E. Wellen, G. Hatzivassiliou, U.M. Sachdeva, T.V. Bui, J.R. Cross, C.B. Thompson, ATP-citrate lyase links cellular metabolism to histone acetylation, Science, 324 (2009) 1076–1080.

[7] S.J. Mentch, M. Mehrmohamadi, L. Huang, X. Liu, D. Gupta, D. Mattocks, P. Gómez Padilla, G. Ables, M.M. Bamman, A.E. Thalacker-Mercer, S.N. Nichenametla, J.W. Locasale, Histone Methylation Dynamics and Gene Regulation Occur through the Sensing of One-Carbon Metabolism, Cell Metab, 22 (2015) 861–873.

[8] M.J. Sadhu, Q. Guan, F. Li, J. Sales-Lee, A.T. Iavarone, M.C. Hammond, W.Z. Cande, J. Rine, Nutritional control of epigenetic processes in yeast and human cells, Genetics, 195 (2013) 831– 844.

[9] N. Shiraki, Y. Shiraki, T. Tsuyama, F. Obata, M. Miura, G. Nagae, H. Aburatani, K. Kume, F. Endo, S. Kume, Methionine metabolism regulates maintenance and differentiation of human pluripotent stem cells, Cell Metab, 19 (2014) 780–794.

[10] N. Shyh-Chang, J.W. Locasale, C.A. Lyssiotis, Y. Zheng, R.Y. Teo, S. Ratanasirintrawoot, J. Zhang, T. Onder, J.J. Unternaehrer, H. Zhu, J.M. Asara, G.Q. Daley, L.C. Cantley, Influence of threonine metabolism on S-adenosylmethionine and histone methylation, Science, 339 (2013) 222–226.

[11] W.-C. Hsieh, B.M. Sutter, H. Ruess, S.D. Barnes, V.S. Malladi, B.P. Tu, Glucose starvation induces a switch in the histone acetylome for activation of gluconeogenic and fat metabolism genes, Mol Cell, 82 (2022) 60–74.e5.

[12] A. Grzenda, G. Lomberk, J.-S. Zhang, R. Urrutia, Sin3: master scaffold and transcriptional corepressor, Biochim Biophys Acta, 1789 (2009) 443–450.

[13] S.M. Cowley, B.M. Iritani, S.M. Mendrysa, T. Xu, P.F. Cheng, J. Yada, H.D. Liggitt, R.N. Eisenman, The mSin3A chromatin-modifying complex is essential for embryogenesis and T-cell development, Mol Cell Biol, 25 (2005) 6990–7004.

[14] G. Pennetta, D. Pauli, The Drosophila Sin3 gene encodes a widely distributed transcription factor essential for embryonic viability, Dev Gene Evol, 208 (1998) 531–536.

[15] L.A. Pile, E.M. Schlag, D.A. Wassarman, The SIN3/RPD3 deacetylase complex is essential for G(2) phase cell cycle progression and regulation of SMRTER corepressor levels, Mol Cell Biol, 22 (2002) 4965–4976.

[16] A. Chaubal, L.A. Pile, Same agent, different messages: insight into transcriptional regulation by SIN3 isoforms, Epigenetics Chromatin, 11 (2018) 17.

[17] D. Kadosh, K. Struhl, Repression by Ume6 involves recruitment of a complex containing Sin3 corepressor and Rpd3 histone deacetylase to target promoters, Cell, 89 (1997) 365–371.

[18] Y.M. Moshkin, T.W. Kan, H. Goodfellow, K. Bezstarosti, R.K. Maeda, M. Pilyugin, F. Karch, S.J. Bray, J.A.A. Demmers, C.P. Verrijzer, Histone chaperones ASF1 and NAP1 differentially modulate removal of active histone marks by LID-RPD3 complexes during NOTCH silencing, Mol Cell, 35 (2009) 782–793.

[19] D. Kadosh, K. Struhl, Targeted Recruitment of the Sin3-Rpd3 Histone Deacetylase Complex Generates a Highly Localized Domain of Repressed Chromatin In Vivo, Mol Cell Biol, 18 (1998) 5121–5127.

[20] P. Gallinari, S. Di Marco, P. Jones, M. Pallaoro, C. Steinkühler, HDACs, histone deacetylation and gene transcription: from molecular biology to cancer therapeutics, Cell Res, 17 (2007) 195–211.

[21] B.E. Bernstein, J.K. Tong, S.L. Schreiber, Genomewide studies of histone deacetylase function in yeast, Proc Natl Acad Sci U S A, 97 (2000) 13708–13713.

[22] J.-H. Dannenberg, G. David, S. Zhong, J. van der Torre, W.H. Wong, R.A. DePinho, mSin3A corepressor regulates diverse transcriptional networks governing normal and neoplastic growth and survival, Genes Dev., 19 (2005) 1581–1595.

[23] A. Gajan, V.L. Barnes, M. Liu, N. Saha, L.A. Pile, The histone demethylase dKDM5/LID interacts with the SIN3 histone deacetylase complex and shares functional similarities with SIN3, Epigenetics & Chromatin, 9 (2016) 4.

[24] L.A. Pile, P.T. Spellman, R.J. Katzenberger, D.A. Wassarman, The SIN3 Deacetylase Complex Represses Genes Encoding Mitochondrial Proteins: IMPLICATIONS FOR THE REGULATION OF ENERGY METABOLISM *, Journal of Biological Chemistry, 278 (2003) 37840–37848.

[25] K. Williams, J. Christensen, M.T. Pedersen, J.V. Johansen, P.A.C. Cloos, J. Rappsilber, K. Helin, TET1 and hydroxymethylcytosine in transcription and DNA methylation fidelity, Nature, 473 (2011) 343– 348.

[26] V.L. Barnes, B.S. Strunk, I. Lee, M. Hüttemann, L.A. Pile, Loss of the SIN3 transcriptional corepressor results in aberrant mitochondrial function, BMC Biochem, 11 (2010) 26.

[27] M. Liu, N. Saha, A. Gajan, N. Saadat, S.V. Gupta, L.A. Pile, A complex interplay between SAM synthetase and the epigenetic regulator SIN3 controls metabolism and transcription, Journal of Biological Chemistry, 295 (2020) 375–389.

[28] M. Liu, V.L. Barnes, L.A. Pile, Disruption of Methionine Metabolism in Drosophila melanogaster Impacts Histone Methylation and Results in Loss of Viability, G3 (Bethesda), 6 (2015) 121–132.

[29] L.A. Pile, D.A. Wassarman, Chromosomal localization links the SIN3–RPD3 complex to the regulation of chromatin condensation, histone acetylation and gene expression, EMBO J, 19 (2000) 6131–6140.

[30] R.L. Aft, F.W. Zhang, D. Gius, Evaluation of 2-deoxy-D-glucose as a chemotherapeutic agent: mechanism of cell death, Br J Cancer, 87 (2002) 805–812.

[31] T. Li, J. Han, L. Jia, X. Hu, L. Chen, Y. Wang, PKM2 coordinates glycolysis with mitochondrial fusion and oxidative phosphorylation, Protein Cell, 10 (2019) 583–594.

[32] Z. Wang, L. Zhang, D. Zhang, R. Sun, Q. Wang, X. Liu, Glycolysis inhibitor 2-deoxy-D-glucose suppresses carcinogen-induced rat hepatocarcinogenesis by restricting cancer cell metabolism, Mol Med Rep, 11 (2015) 1917–1924.

[33] N. Saha, M. Liu, A. Gajan, L.A. Pile, Genome-wide studies reveal novel and distinct biological pathways regulated by SIN3 isoforms, BMC Genomics, 17 (2016) 111.

[34] Y. Tsukada, J. Fang, H. Erdjument-Bromage, M.E. Warren, C.H. Borchers, P. Tempst, Y. Zhang, Histone demethylation by a family of JmjC domain-containing proteins, Nature, 439 (2006) 811– 816.

[35] C.F. Bennett, C.T. Ronayne, P. Puigserver, Targeting adaptive cellular responses to mitochondrial bioenergetic deficiencies in human disease, FEBS J, 289 (2022) 6969–6993.

[36] D.H. Jang, M. Kelly, K. Hardy, D.S. Lambert, F.S. Shofer, D.M. Eckmann, A preliminary study in the alterations of mitochondrial respiration in patients with carbon monoxide poisoning measured in blood cells, Clin Toxicol (Phila), 55 (2017) 579–584.

[37] A.S. Divakaruni, M.D. Brand, The Regulation and Physiology of Mitochondrial Proton Leak, Physiology, 26 (2011) 192–205.

[38] S.A. Mookerjee, D.G. Nicholls, M.D. Brand, Determining Maximum Glycolytic Capacity Using Extracellular Flux Measurements, PLOS ONE, 11 (2016) e0152016.

[39] M. Bucher, L. Kadam, K. Ahuna, L. Myatt, Differences in Glycolysis and Mitochondrial Respiration between Cytotrophoblast and Syncytiotrophoblast In-Vitro: Evidence for Sexual Dimorphism, Int J Mol Sci, 22 (2021) 10875.

[40] M.M. Spain, J.A. Caruso, A. Swaminathan, L.A. Pile, Drosophila SIN3 Isoforms Interact with Distinct Proteins and Have Unique Biological Functions, J Biol Chem, 285 (2010) 27457–27467.

[41] J. Zecha, S. Satpathy, T. Kanashova, S.C. Avanessian, M.H. Kane, K.R. Clauser, P. Mertins, S.A. Carr, B. Kuster, TMT Labeling for the Masses: A Robust and Cost-efficient, In-solution Labeling Approach, Mol Cell Proteomics, 18 (2019) 1468–1478.

[42] K.T. Smith, M.E. Sardiu, S.A. Martin-Brown, C. Seidel, A. Mushegian, R. Egidy, L. Florens, M.P. Washburn, J.L. Workman, Human family with sequence similarity 60 member A (FAM60A) protein: a new subunit of the Sin3 deacetylase complex, Mol Cell Proteomics, 11 (2012) 1815–1828.

[43] G. Streubel, D.J. Fitzpatrick, G. Oliviero, A. Scelfo, B. Moran, S. Das, N. Munawar, A. Watson, K. Wynne, G.L. Negri, E.T. Dillon, S. Jammula, K. Hokamp, D.P. O’Connor, D. Pasini, G. Cagney, A.P. Bracken, Fam60a defines a variant Sin3a-Hdac complex in embryonic stem cells required for self-renewal, EMBO J, 36 (2017) 2216–2232.

[44] I. Soukar, A. Mitra, L.A. Pile, Analysis of the chromatin landscape and RNA polymerase II binding at SIN3-regulated genes, Biol Open, 12 (2023) bio060026.

[45] I. Schneider, Cell lines derived from late embryonic stages of Drosophila melanogaster, J Embryol Exp Morphol, 27 (1972) 353–365.

[46] L.S. Gramates, J. Agapite, H. Attrill, B.R. Calvi, M.A. Crosby, G. dos Santos, J.L. Goodman, D. Goutte-Gattat, V.K. Jenkins, T. Kaufman, A. Larkin, B.B. Matthews, G. Millburn, V.B. Strelets, the FlyBase Consortium, FlyBase: a guided tour of highlighted features, Genetics, 220 (2022) iyac035.

[47] S.X. Ge, D. Jung, R. Yao, ShinyGO: a graphical gene-set enrichment tool for animals and plants, Bioinformatics, 36 (2020) 2628–2629.

[48] W. Luo, C. Brouwer, Pathview: an R/Bioconductor package for pathway-based data integration and visualization, Bioinformatics, 29 (2013) 1830–1831.

[49] A. Mitra, L. Vo, I. Soukar, A. Chaubal, M.L. Greenberg, L.A. Pile, Isoforms of the transcriptional cofactor SIN3 differentially regulate genes necessary for energy metabolism and cell survival, (2022) 2021.12.31.474661.

[50] A. Mitra, A.-M. Raicu, S.L. Hickey, L.A. Pile, D.N. Arnosti, Soft repression: Subtle transcriptional regulation with global impact, Bioessays, 43 (2021) e2000231.

[51] F. Buttgereit, M.D. Brand, A hierarchy of ATP-consuming processes in mammalian cells, Biochem J, 312 ( Pt 1) (1995) 163–167.

[52] R. Vaid, J. Wen, M. Mannervik, Release of promoter–proximal paused Pol II in response to histone deacetylase inhibition, Nucleic Acids Research, 48 (2020) 4877–4890.

